# ’Simple Tidy GeneCoEx’: a gene co-expression analysis workflow powered by tidyverse and graph-based clustering in R

**DOI:** 10.1101/2022.11.11.516131

**Authors:** Chenxin Li, C. Robin Buell

## Abstract

Gene co-expression analysis is an effective method to detect groups (or modules) of co-expressed genes that display similar expression patterns, which may function in the same biological processes. Here, we present ‘Simple Tidy GeneCoEx’, a gene co-expression analysis workflow written in the R programming language. The workflow is highly customizable across multiple stages of the pipeline including gene selection, edge selection, clustering resolution, and data visualization. Powered by the tidyverse package ecosystem and network analysis functions provided by the igraph package, the workflow detects gene co-expression modules whose members are highly interconnected. Step-by-step instructions with two use case examples as well as source code are available at https://github.com/cxli233/SimpleTidy_GeneCoEx.

**Core Ideas:**

- An R-based workflow that performs gene co-expression analysis was developed.
- The workflow is based on tidyverse packages and graph theory.
- The workflow is highly customizable, detects tight gene co-expression modules, and generates publication quality figures.
- Two plant gene expression datasets were used to benchmark the workflow.

## 1. Introduction

Transcriptomic analyses have become routine for studying plant biology. A challenge for plant biologists is interpreting omics data to derive biological insights. A valuable and powerful tool for gene expression analyses is gene co-expression. When multiple treatments (time points, developmental stages, cell types, genotypes, and perturbations) are included in a gene expression study, it is possible to detect groups of genes, or gene co-expression modules, with similar expression profiles across a range of treatment conditions or through a developmental timepoints. Under the ‘guilt-by-association’ assumption, genes with expression patterns similar to previously characterized genes with known roles in a biological process (bait genes) are deduced to function in the same biological process. In addition, candidate genes of interest can be detected in modules with interesting expression patterns, which can then be subjected to further forward or reverse genetics studies. Gene co-expression analyses have been successfully applied to identify genes implicated in development, stress responses, primary metabolism, and specialized metabolism across a wide range of plant species including crops and medicinal plants (Burlat et al. 2004; Anderson et al. 2017; Gomez-Cano et al. 2022; Moghaddam et al. 2021).

Due to its general ease of use, open-source nature, and availability of general and domain-specific packages, the R programming language for statistical computing has become the programming language of choice for gene expression and computational biology analyses (Tippmann 2015). Within the R programming environment, the tidyverse ecosystem is a collection of packages built upon a common programming style, grammar, and data structures (Wickham et al. 2019). A key underlying concept of the tidyverse ecosystem is ‘tidy data frames’ which are data frames with observations as rows and variables as columns. The ‘tidy’ nature of data frames greatly facilitates grouping, filtering, joining, reshaping, summarizing, and visualizing data using tidyverse functions. Since gene expression matrixes are also tabular in nature, gene co-expression analyses can be done in a tidyverse-compatible manner. Tidy data frames can be seamlessly integrated with igraph (Csárdi and Nepusz 2006), a powerful network analysis package in R, as igraph contains methods that converts data frames into network objects. In graph theory, a network is considered a graph, a mathematical structure used to model pairwise relationships. Thus, the pairwise correlations among genes can be modeled by a graph in which genes are nodes and correlations are edges. Further, gene co-expression modules can be detected by graph-based clustering. Here, we developed a gene co-expression workflow ‘Simple Tidy GeneCoEx’ using tidyverse and igraph functions. The workflow is highly customizable across multiple stages of the pipeline, including gene selection, edge selection, clustering resolution, and data visualization. Step-by-step instructions for two benchmarked use cases are available at https://github.com/cxli233/SimpleTidy_Gen-eCoEx.

## 2. Methods

### 2.1 Overview

A straightforward pipeline was designed with plant molecular biologists and geneticists in mind: (i) import gene expression matrix, (ii) filter for genes that are expressed, exhibit high variance, and/or high F statistics, (iii) produce correlation matrix and filter edges, (iv) detect gene co-expression modules, and (vi) plot/export results. The workflow is executed by calling tidyverse (Wickham et al. 2019) and igraph (Csárdi and Nepusz 2006) functions.

### 2.2 Test Datasets

The workflow has been tested on two distinct datasets: tomato fruit developmental series (Shinozaki et al. 2018) and tepary bean heat stress time course (Moghaddam et al. 2021). The tomato fruit developmental series dataset contains six hand dissected tissues and five laser capture micro-dissected (LCM) tissues across 11 developmental stages, ranging from anthesis to red ripe (i.e., fully ripe tomato fruits). For simplicity of demonstration, only hand dissected samples (*n* = 84 unique tissue by developmental stage combinations) were analyzed by this workflow, as it has been noted that the LCM samples were lower input, constructed by a different library preparation kit, and had globally distinct expression pattern relative to hand dissected samples (Shinozaki et al. 2018). The tepary bean stress time course experiment contains two treatments (control vs. heat) and five time points over a 24-hr period (1, 3, 6, 12, and 24 hours post stress), an experiment with a strong diurnal component (Moghaddam et al. 2021). All treatment by time point combinations (*n* = 10 combinations) were used in the test analyses. These datasets were chosen because of their multifactorial experimental designs and distinct biological questions (development and stress) that were investigated.

### 2.3 Required inputs

The workflow requires three inputs: (1) gene expression matrix, (2) library metadata, and (3) bait genes. A variety of software can be used to generate gene expression matrices, such as Cufflinks (Trapnell et al. 2012), kallisto (Bray et al. 2016), and STAR (Dobin et al. 2013). The required format is that each row is a gene, and each column is a biological sample. Values in the gene coexpression matrix should be depth and normalized gene expression estimates, in units of transcripts per million (TPM) or fragments per kilobase of exon model per million mapped fragments (FPKM). A metadata table is required for the workflow, in which each row corresponds to a sample (i.e., sequencing library), and columns correspond to biological and technical aspects of the libraries. Finally, a table of bait genes is used to guide the pipeline, since oftentimes users have prior knowledge of genes involved in the biological processes being studied. The required format is that each row is a gene. Additional information about bait genes such as functional annotations and genomic locations can be recorded as columns in the bait gene table. Before starting the workflow, exploratory analyses, such as principal component analysis (PCA) are encouraged to examine the major drivers of variance among samples.

### 2.4 Gene selection

Gene selection prior to co-expression analysis is optional. However, since the workflow constructs all pairwise correlations among genes, the number of correlations scales with the square of number of genes in the analyses. Thus, pre-filtering genes can significantly speed up the workflow. Gene selection can be performed using one or more of the following methods: expression threshold, variance threshold, and F statistics threshold.

Gene selection based on expression value is the most conceptually simple. It asks if a given gene is expressed among the samples being analyzed, given an expression threshold *E* and prevalence threshold *N*_*P*_. A simple method is to subset genes with expression values > *E* in at least *N*_*P*_ libraries, where the values for *E* and *N*_*P*_ can be determined by the users based on the dataset. A recommendation for selecting a prevalence threshold is to use the lowest level of replication across treatments. For example, across all treatments in a study, if the treatment with the least number of biological replicates has three replicates, then a recommended prevalence cutoff is *N*_*P*_ = 3.

More involved methods of gene selection are based on biological variance and F statistics. For gene selection based on biological variances, the underlying assumption is that genes distinctly expressed in one or more treatments have higher biological variances than genes expressed at similar levels across all treatments. In this workflow, technical variation is reduced by first averaging replicates to the level of the treatments. To reduce the bias towards highly expressed the genes, pre-filtering high variance genes is done by first log-transforming the expression value, then averaging replicates up to the level of treatments, and finally selecting high variance genes at the log-transformed scale. Biological variance of bait genes can be used to determine the variance threshold. For example, if user-selected bait genes are ranked among the highest 5000 variable genes, then the top 5000 variable genes can be selected for downstream analyses (Fig. 1a, data from (Shinozaki et al. 2018)).

**Fig. 1.**
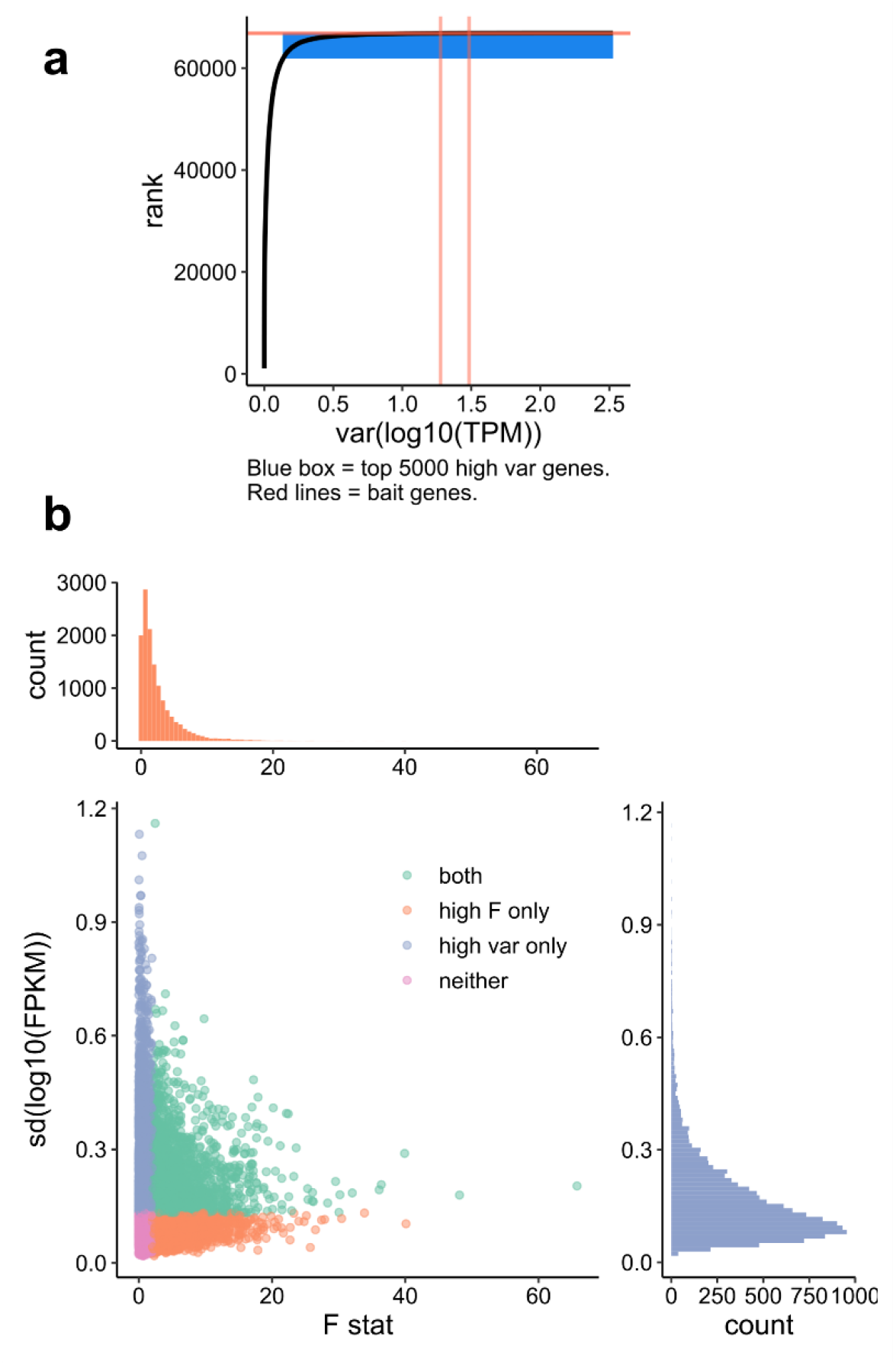
Gene pre-filtering using biological variance and F statistics. (a) Rank vs. value plot for transcripts (data from Shinozaki et al., 2018). Blue box includes top 5000 variable genes, and orange lines correspond to two user-provided bait genes (Solly.M82.10G020850.1 and Solly.M82.03G005440.1). In this analyses, the top 5000 variable transcripts were used for downstream analyses. (b) Scatter plot showing standard deviation (sd) and F statistics of expressed genes (data from Moghaddam et al. 2021). In this case, filtering for high variance or high F statistics (*F* > 2) do not select for the same set of genes. In this analysis, the union of high variance and high F genes were used for downstream analyses.

An alternative gene selection method to biological variance is the F statistics, which detects genes whose expression levels are changing across treatments. The F statistics is computed by first fitting a linear model for each gene:

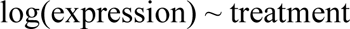

The dependent variable is log-transformed to reduce the heteroscedasticity and mean-error relationship associated with gene expression data. If the experiment is multifactorial in nature, then users have the option to fit the linear model with the single factor accounting for the most variation in the dataset, or the interactions among two or more factors. Depending on the independent variable(s) in the model, the F statistics reflect if a gene is changing expression across a single factor or across the combinations of multiple factors. The F statistics are then calculated by ANOVA. After the F statistics are computed for each gene, genes can be filtered by the F statistics values. We discourage the use of p value for this gene selection method since most gene expression experiments have low levels of replication (typically *n* = 3). As a result, selecting F statistics using p value is overly conservative. Instead, we recommend an F statistics cutoff between 2 to 3. Depending on the model, high biological variance or high F statistics are not mutually exclusive, nor do they select for the same set of genes (Fig. 1b, data from (Moghaddam et al. 2021)). Depending on the biological questions of interest, high variance genes, high F statistics genes, or the union can be used for downstream analyses.

### 2.5 Edge selection

Gene selection produces the nodes of the graph object for downstream network analyses. To construct edges of the network, the workflow uses pairwise gene correlation on standardized log-transformed expression values (z scores of log-transformed expression values). The correlation matrix contains the Pearson correlation coefficient *r* of all pairwise correlations. A p value can be computed from each correlation coefficient, which are then adjusted for multiple comparisons using the Benjamini-Hochberg procedure (Benjamini and Hochberg 1995). However, we encourage users to derive an *r* cutoff based on empirical observations of bait genes instead of using adjusted p values alone, since p value is affected by both *r* and degrees of freedom. Experiments with larger number of treatments and thus higher degrees of freedom produce smaller p values given the same *r* value. As a result, in experiments with large numbers of treatments, selecting an *r* cutoff based solely on p values will be too non-stringent. Instead, prior knowledge regarding bait genes can be used to guide edge selection. For example, users can examine the correlation between two bait genes known to be co-expressed and select an *r* cutoff accordingly (Fig. 2, data from (Shinozaki et al. 2018)). Alternatively, edge selection can be done using mutual ranks (Wisecaver et al. 2017; Obayashi and Kinoshita 2009).

**Fig. 2.**
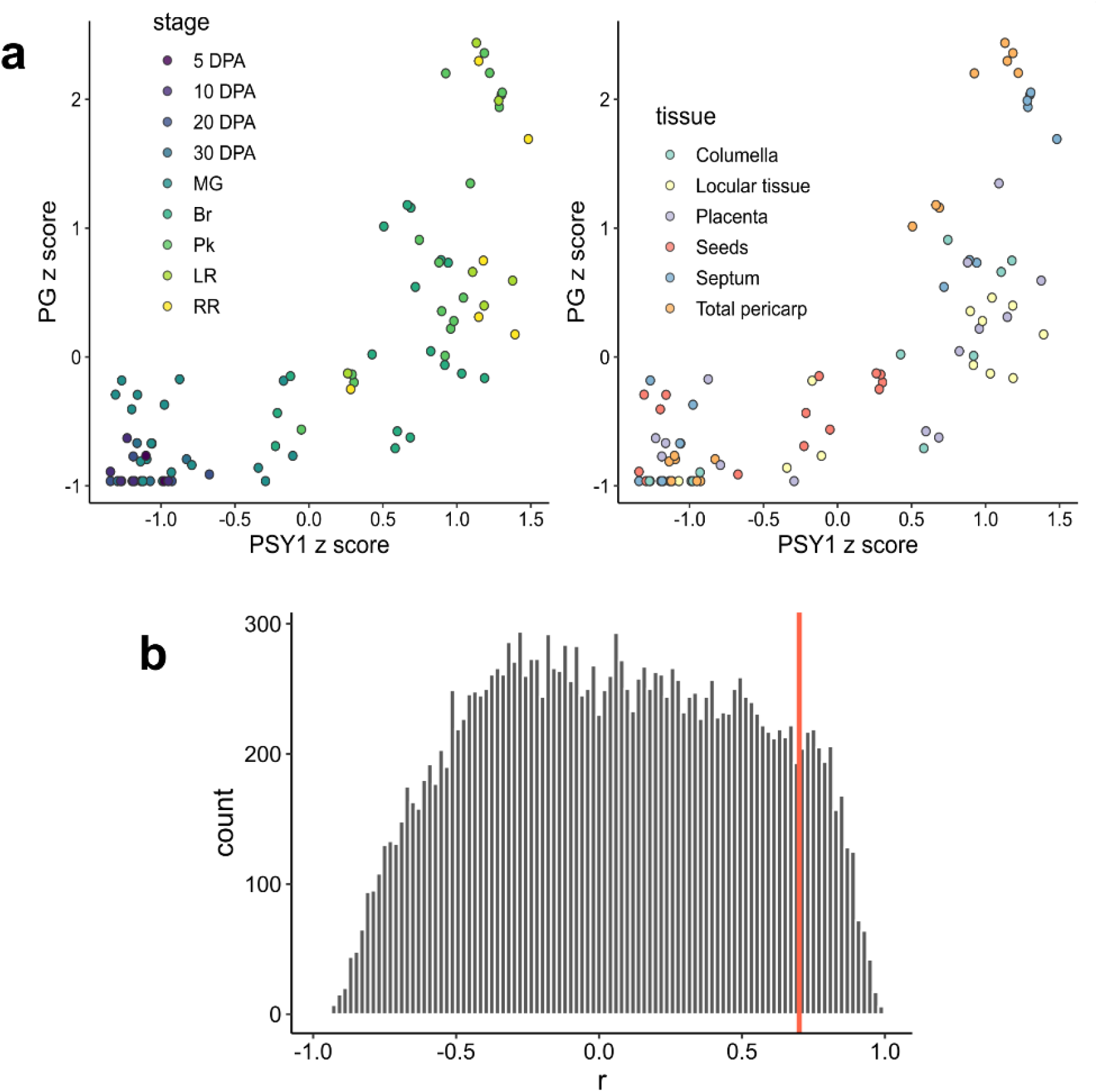
Edge selection using bait genes. (a) Scatter plots showing standardized z scores of *PSY1* and *PG*, two genes previously known to be co-expressed (data from Shinozaki et al., 2018), *r* = 0.75. DPA: days post anthesis. MG: mature green. Br: breaker. Pk: pink. LR: light red. RR: red ripe. (b) Histogram showing distribution of correlation coefficient *r*. Based on correlation coefficient of known co-expressed genes (shown in **a**), the cutoff is chosen at *r* > 0.7 (red line), beyond which the histogram drops off rapidly.

### 2.6 Construction of the network object and graph-based clustering

The nodes (genes) and edges (correlations) are passed onto the ‘*graph_frome_data_frame()*’ function of igraph to generate the network object for graph-based clustering. Gene co-expression modules are then detected using the Leiden algorithm (Traag, Waltman, and van Eck 2019), which detects modules whose members are highly interconnected. The Leiden algorithm is implemented using the ‘*cluster_leiden()*’ function within the igraph package. A critical parameter for module detection is resolution, which needs to be optimized for each experiment. Too low of a resolution forces genes with different expression patterns into a single module, whereas too high of a resolution leads to many genes not contained in a module. The resolution parameter can be optimized by testing a range of resolution values and monitoring the number of modules with 5 or more genes, as well as the number of genes contained in modules with 5 or more genes. The minimum module size 5 is chosen arbitrarily, but generally, higher resolution leads to more modules but less genes contained in large modules (Fig 3).

**Fig. 3.**
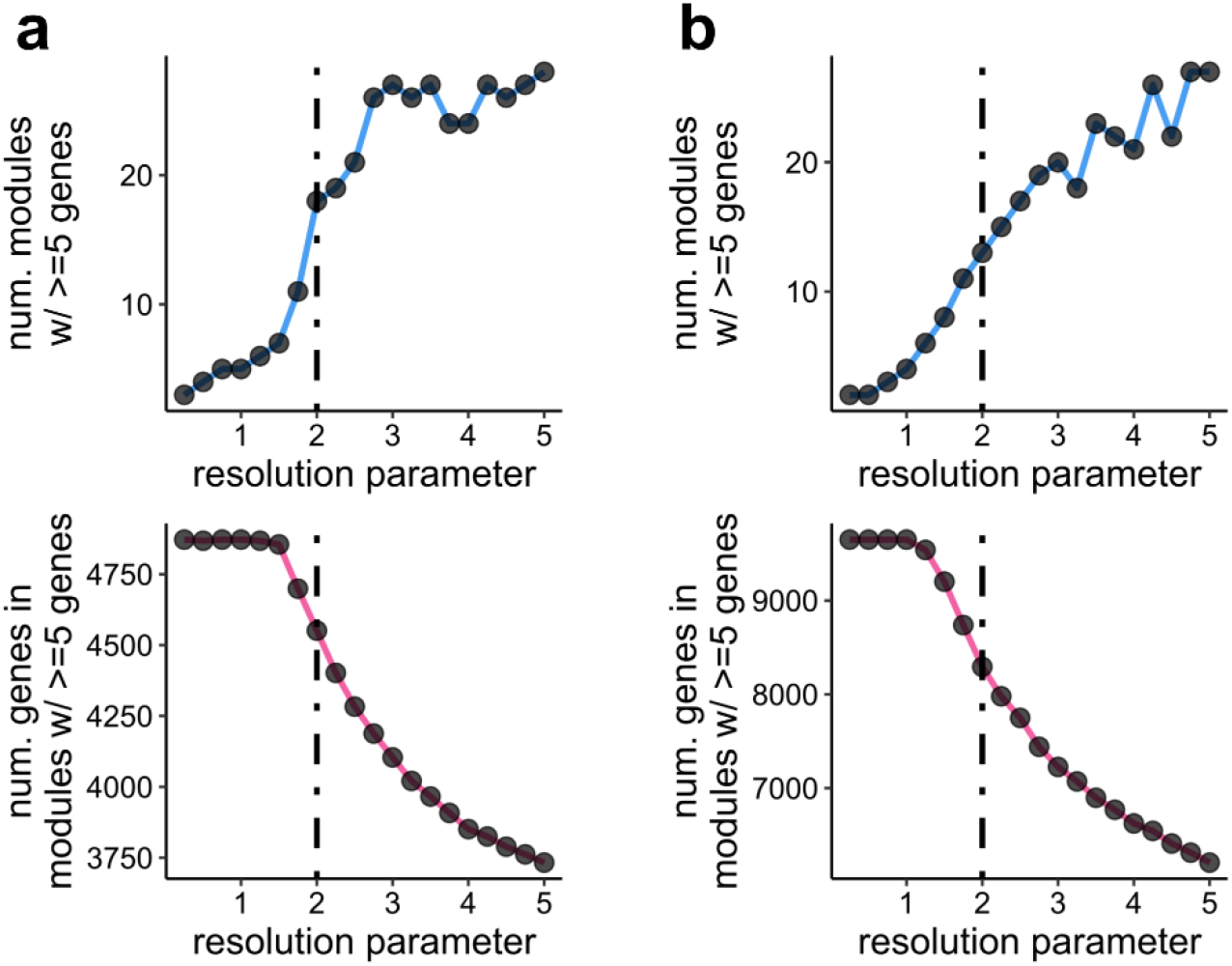
Resolution for graph-based clustering. (a) Tradeoff between module number and genes retained (data from Shinozaki et al., 2018). (b) Tradeoff between module number and genes retained (data from Moghaddam et al. 2021). Dotted lines represent a resolution of 2, a comprise between two the performance indexes.

## 3. Results

### 3.1 Data visualization

From the gene co-expression modules detected by this workflow, a few data visualization options are available, such as heatmap and line graphs (Fig. 4). For heatmaps, the workflow reorders rows and columns based on module peak expression. The workflow was tested on two distinct use cases: tomato fruit developmental series (Shinozaki et al. 2018) (Fig. 4a) and tepary bean heat stress time course (Moghaddam et al. 2021) (Fig. 4b). The workflow can detect gene co-expression modules that are highly expressed in early fruit development (e.g., Fig. 4a, Module 137) and fruit ripening (Fig. 4a, Module 9), as well as tissue specific modules (Fig. 4a, Module 8, a seed specific module). The workflow appears to perform well for experiments with a strong diurnal component, as indicated by the detection of modules that appeared to cycle within a 24-hr period (Moghaddam et al. 2021) (Fig. 4b, Module 7), in addition to stress-responsive modules (Fig. 4b, Modules 3 and 9). The workflow also provides methods for candidate gene identification using module membership, as well as querying direct neighbors to bait genes using the ‘*neighbors()*’ function within igraph. Expression values of candidate genes (in the original scale or log-transformed scale) as well as dispersion among replicates can be visualized (Fig. 4c).

**Fig. 4.**
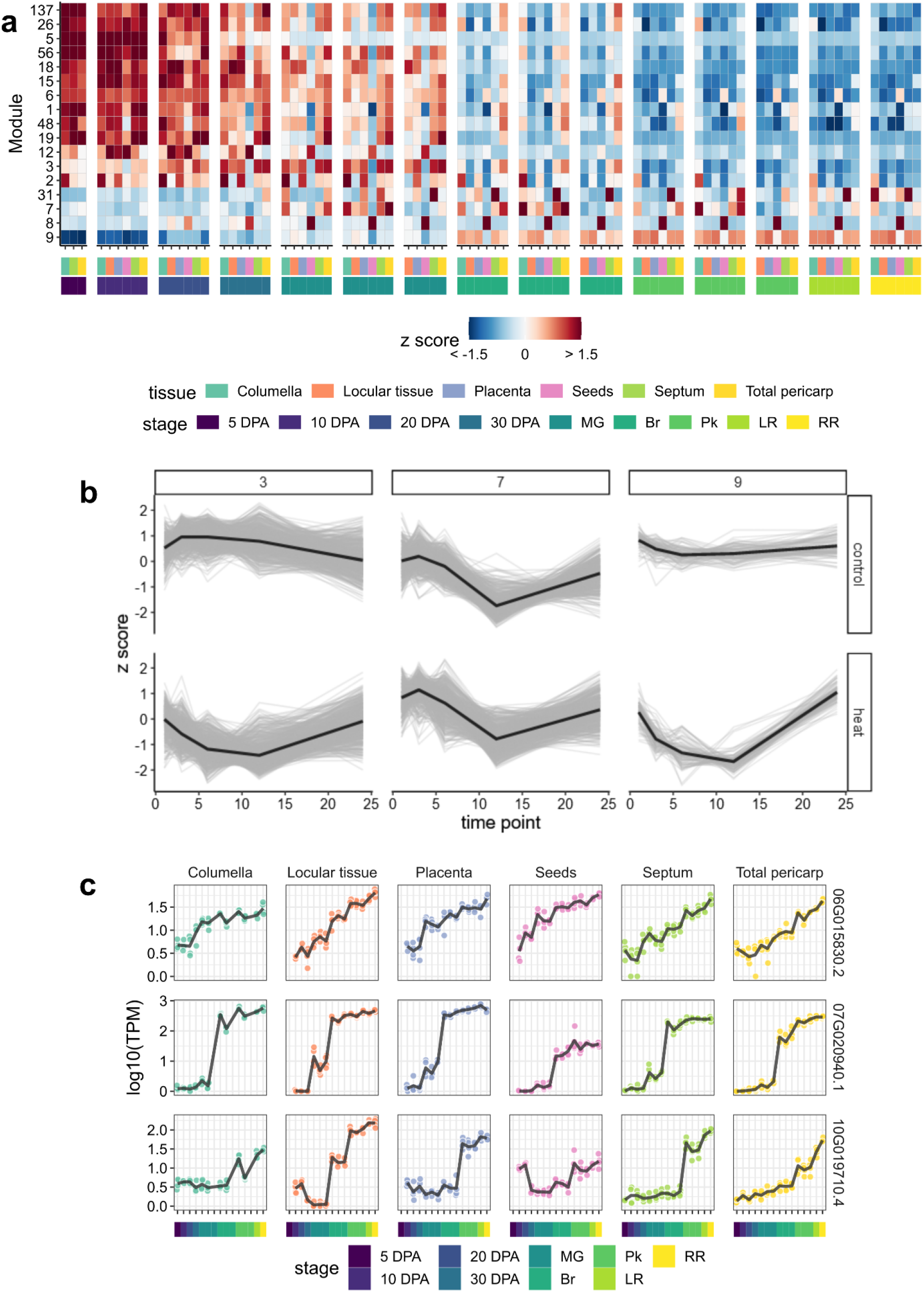
Heatmap and line graph visualization for gene co-expression modules. (a) Heatmap for gene co-expression modules (data from Shinozaki et al., 2018). (b) Line graphs for gene co-expression modules (data from Moghaddam et al. 2021). Thin grey lines represent individual genes. Black lines represent the average expression pattern of the module. (c) Line graphs showing exemplar candidate genes based on module membership (Module 9 in **a**) as well as network neighborhood to bait genes (data from Shinozaki et al., 2018).

### 3.4 Benchmarking against WGCNA

We benchmarked our ‘Simple Tidy GeneCoEx’ method against Weighted Gene Coexpression Network Analysis (WGCNA), a widely accepted gene co-expression analysis package (Langfelder and Horvath 2008) using both use cases (tomato fruit development and tepary bean stress time course) (Shinozaki et al. 2018; Moghaddam et al. 2021). We found that both methods can detect treatment-specific/enriched gene co-expression modules. While there was a lack of a one-to-one correspondence between modules detected by the two methods, we detected groups of modules with similar expression patterns. For example, the “plum1” Module detected by WGCNA is highly correlated with Module 9 detected by this workflow; both peaked at the red ripe stage of tomato fruit development (Fig. 5). Analysis of module membership revealed the equivalence of a subset of modules detected by either method (Fig. 6). In some cases, the two methods detected modules that share practically the same membership, while in other cases, a large module detected by one method is split into multiple smaller modules that have similar expression patterns by the other method.

**Fig. 5.**
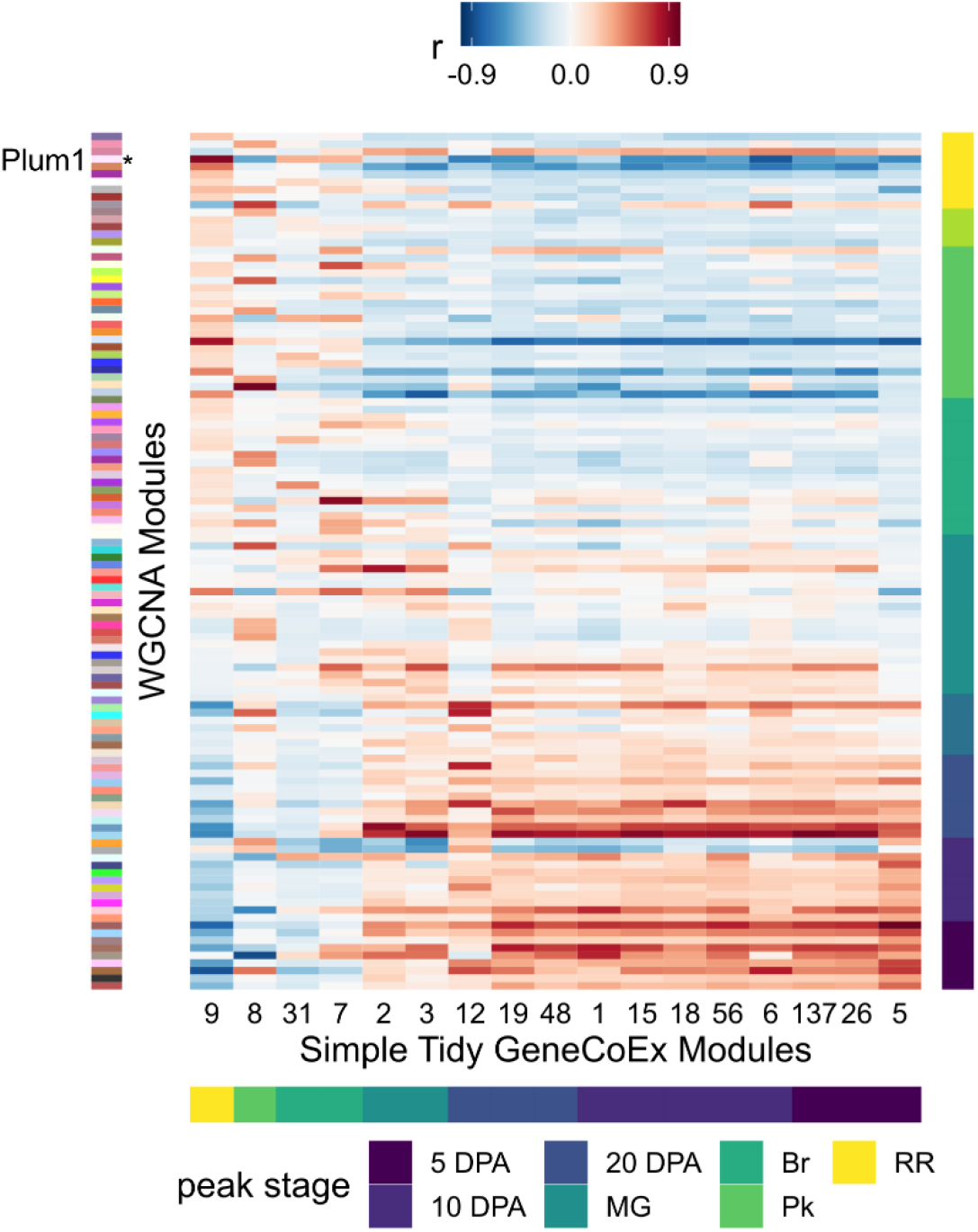
Module correspondence between WGCNA and ‘Simple Tidy Gene CoEx’. Rows are gene co-expression modules detected by WGCNA, annotated by the color strip on the left. Columns are modules detected by ‘Simple Tidy GeneCoex’. Color strips at the bottom or on the right annotate the module peak expression. Heatmap colors indicate correlation coefficient (*r*).

**Fig. 6.**
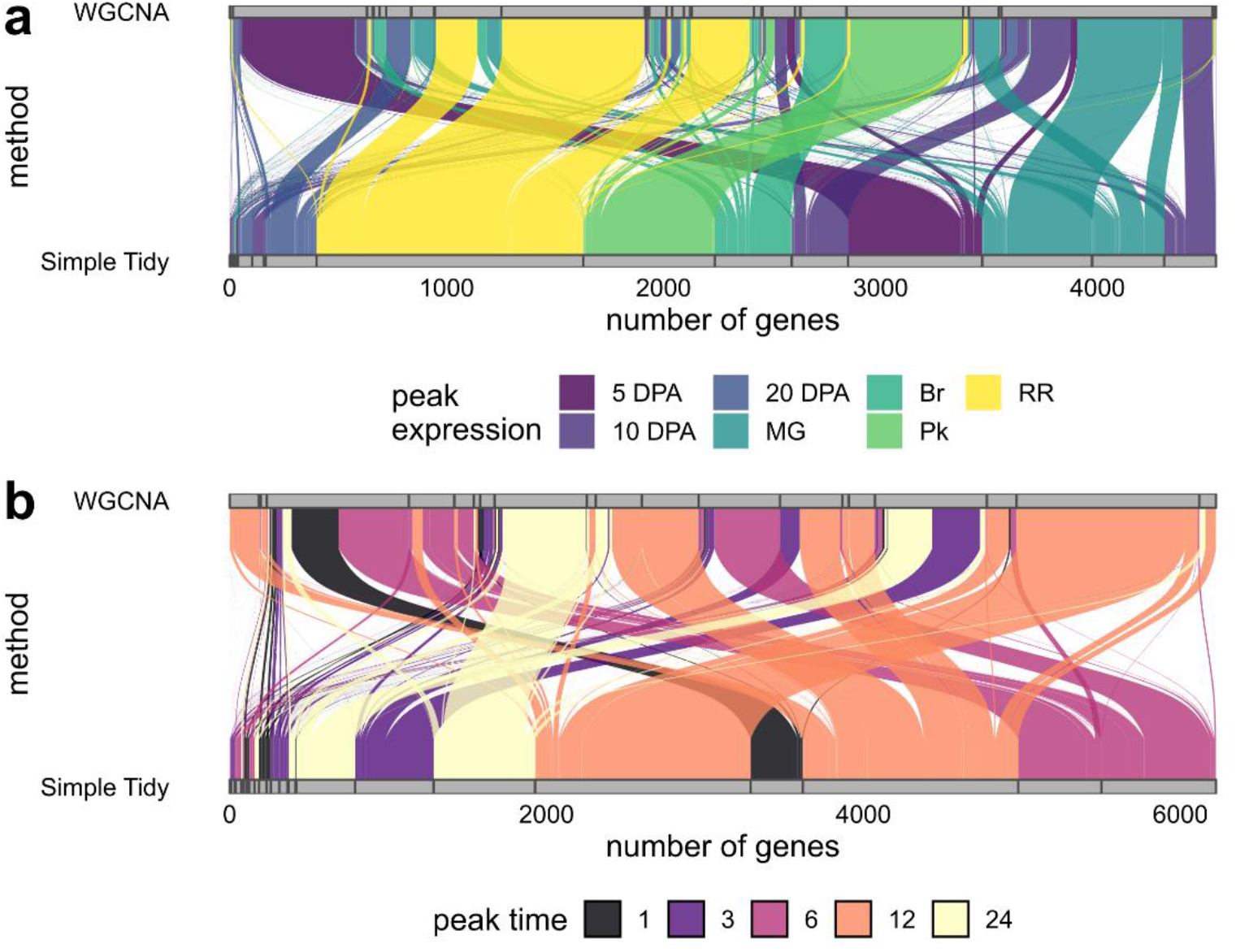
Membership analyses between two gene co-expression methods, visualized by alluvial plots. Horizontal grey bars represent gene co-expression modules. Blocks of colored curves represent shared membership. (a) data from Shinozaki et al. (2018). (b) data from Moghaddam et al. (2021).

### 3.3 Module tightness

To evaluate and compare the quality or tightness of gene co-expression modules detected by either method, we computed the squared error loss for each module, which is defined as:

For gene *i* and treatment *j* in module *m*, the mean sum of square of such a module, i.e., *msq*_*m*_, is computed by:

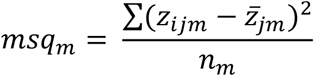

where *z*_*ij*_ is the z score of each gene at each treatment, 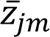 is the average z score across all genes in the module at each treatment, and *n*_*m*_ is the total number of genes in each module, such that the sum of squares is normalized to the number of genes in each module.

We computed *msq*_*m*_ for each module detected by WGCNA or ‘Simple Tidy GeneCoEx’ and found that consistently for both use cases, the ‘Simple Tidy GeneCoEx’ workflow detected modules with lower squared loss error (Fig. 7). For the Shinozaki et al. (2018) data, there was a ∼45% reduction in *msq*_*m*_ using ‘Simple Tidy GeneCoEx’ relative to WGCNA (*P* = 3.6 × 10^−8^, Wilcoxon Rank Sum Test). The association between *msq*_*m*_ and module size (number of genes in modules) was weak (*r* = 0.17), suggesting the higher *msq*_*m*_ values for WGCNA modules is not due to insufficient clustering resolution (Fig. 7a). For data from Moghaddam et al. (2021) data, we saw a ∼40% reduction in *msq*_*m*_ using ‘Simple Tidy GeneCoEx’ relative to WGCNA (*P* = 3.1 × 10^−5^, Wilcoxon Rank Sum Test). We also observed a mild association between module size and *msq*_*m*_ (*r* = 0.526), suggesting both methods may benefit from a higher clustering resolution (Fig. 7b). However, after controlling for module size using a mixed effect linear model with module size as a random effect covariate, on average, the ‘Simple Tidy GeneCoEx’ workflow returned lower *msq*_*m*_ values (estimate = -0.939, 95% confidence interval = [-1.6, -0.276], *F* = 8.6, *P* = 0.0067, ANCOVA). Taken together, the ‘Simple Tidy GeneCoEx’ workflow detects gene co-expression modules that are tighter than those detected by WGCNA.

**Fig. 7.**
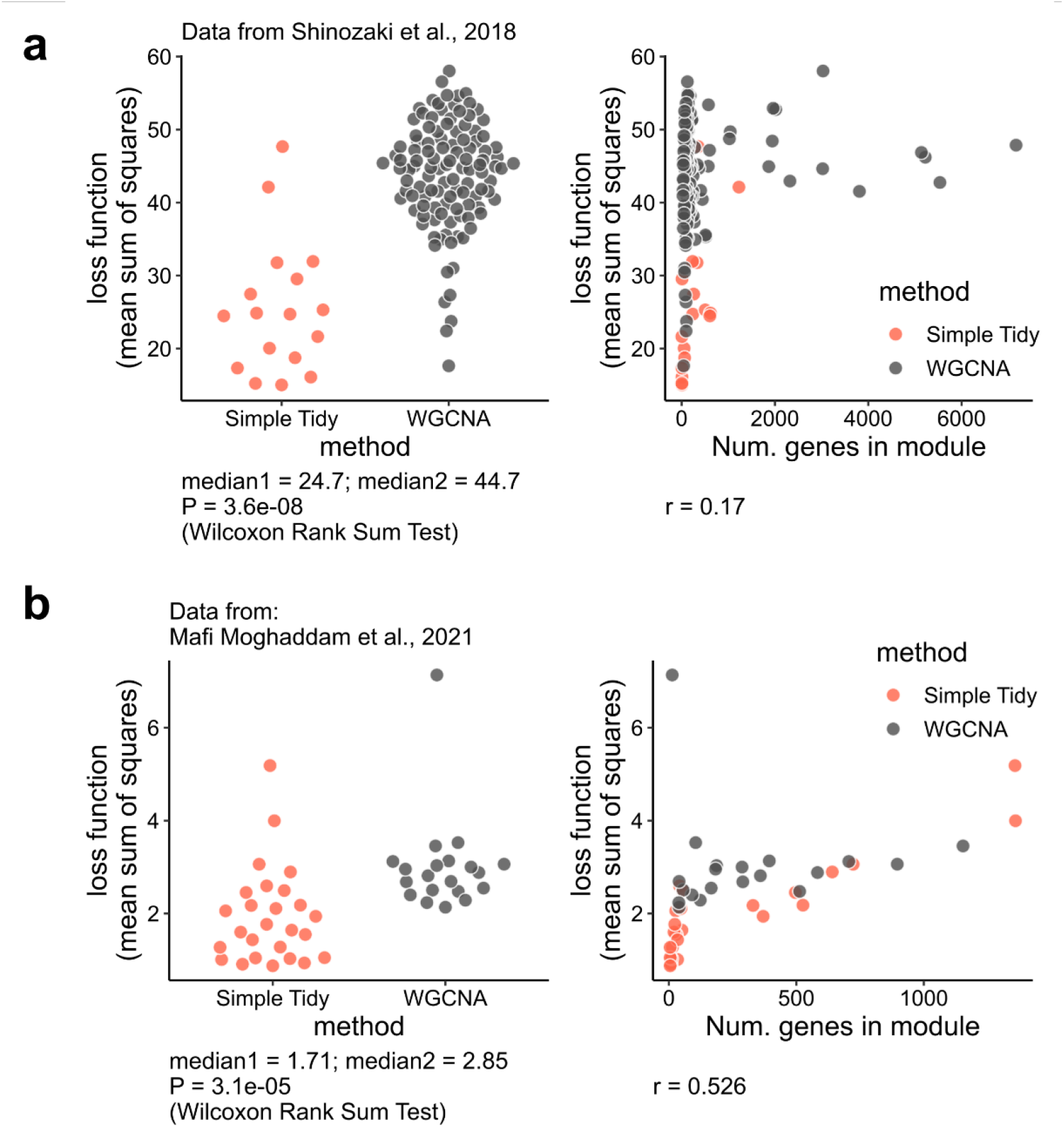
Quantification of module tightness. Each data pot is a module, color coded by the gene co-expression method. (a) data from Shinozaki et al. (2018). (b) data from Moghaddam et al. (2021).

## 4. Discussion

Here, we present a simple, highly customizable co-expression analysis workflow in R powered by tidyverse and igraph functions. The workflow has been tested on two distinct gene expression studies (Shinozaki et al. 2018; Moghaddam et al. 2021), one focused on development and one focused on a diurnal time course following heat stress. The workflow is applicable to other gene expression studies such as single cell RNA-seq experiments. In a recent study, we applied this workflow to detected co-expression modules enriched in specific cell types, which were used to discover candidate genes in a biosynthetic pathway for complex plant natural products (Li et al. 2022). The method has been benchmarked against WGCNA, a widely accepted gene co-expression package. We found that across two distinct use cases, the ‘Simple Tidy GeneCoEx’ method detects modules that are, on average, tighter than those detected by WGCNA. A potential reason underlying the differences in module tightness might be due to the module detection methods. By default, WGCNA uses hierarchical clustering followed by tree cutting to detect modules (Langfelder, Zhang, and Horvath 2008). In contrast, ‘Simple Tidy GeneCoEx’ uses the Leiden algorithm to detect modules, which returns modules that are highly interconnected (Traag, Waltman, and van Eck 2019).

## Data availability

Gene expression matrix for Shinozaki et al. (2018) are available at Zenodo: https://zenodo.org/rec-ord/7117357. Gene expression matrix for Moghaddam et al. (2021) are available at GitHub: https://github.com/cxli233/SimpleTidy_GeneCoEx/tree/main/Data/Moghaddam2022_data. Step-by-step instructions for the workflow and source code are available at GitHub https://github.com/cxli233/SimpleTidy_GeneCoEx, and stable release of source code are available at Zenodo: https://zenodo.org/record/7182680.

## Author Contributions

CL conceived the study, developed the pipeline, prepared figures, and wrote the manuscript with input from CRB.

## Acknowledgements

We thank Dr. Daniel Kliebenstein and John Hamilton for discussions regarding the pipeline. We also thank Dr. Natalie Deans for producing the gene expression matrix from data published by Shinozaki et al. (2018) using kallisto. We thank Drs. Mitchell Feldmann, Valerio Hoyos-Villegas, Seth Murray, and Jason Wallace for discussions regarding mixed effect linear models. Funds from the Georgia Research Alliance and the University of Georgia to CRB supported this work.

## Abbreviations

ANCOVA: analysis of covariance
ANOVA: analysis of variance
FPKM: fragments per kilobase exon model per million mapped fragments
LCM: laser capture micro-dissection
msq: mean sum of squares
PCA: principal component analysis
sd: standard deviation
TPM: transcripts per million
WGCNA: weighted gene co-expression network analysis

## Reference

Anderson, Sarah N., Cameron S. Johnson, Joshua Chesnut, Daniel S. Jones, Imtiyaz Khanday, Margaret Woodhouse, Chenxin Li, Liza J. Conrad, Scott D. Russell, and Venkatesan Sundaresan. 2017. “The Zygotic Transition Is Initiated in Unicellular Plant Zygotes with Asymmetric Activation of Parental Genomes.” Developmental Cell 43 (3): 349-358.e4. https://doi.org/10.1016/j.devcel.2017.10.005.

Benjamini, Yoav, and Yosef Hochberg. 1995. “Controlling the False Discovery Rate: A Practical and Powerful Approach to Multiple Testing.” Journal of the Royal Statistical Society. Series B 57 (1): 289–300.

Bray, Nicolas L, Harold Pimentel, Páll Melsted, and Lior Pachter. 2016. “Near-Optimal Probabilistic RNA-Seq Quantification.” Nature Biotechnology 34 (5): 525–27. https://doi.org/10.1038/nbt.3519.

Burlat, Vincent, Audrey Oudin, Martine Courtois, Marc Rideau, and Benoit St-Pierre. 2004. “Co-Expression of Three MEP Pathway Genes and Geraniol 10-Hydroxylase in Internal Phloem Parenchyma of Catharanthus Roseus Implicates Multicellular Translocation of Intermediates during the Biosynthesis of Monoterpene Indole Alkaloids and Isoprenoid-Derived Primary Metabolites.” The Plant Journal 38 (1): 131–41. https://doi.org/10.1111/j.1365-313X.2004.02030.x.

Csárdi, Gábor, and Tamás Nepusz. 2006. “The Igraph Software Package for Complex Network Research.” InterJournal, Complex Systems, 1695. https://igraph.org.

Dobin, Alexander, Carrie A. Davis, Felix Schlesinger, Jorg Drenkow, Chris Zaleski, Sonali Jha, Philippe Batut, Mark Chaisson, and Thomas R. Gingeras. 2013. “STAR: Ultrafast Universal RNA-Seq Aligner.” Bioinformatics 29 (1): 15–21. https://doi.org/10.1093/bioinformatics/bts635.

Gomez-Cano, Fabio, Yi-Hsuan Chu, Mariel Cruz-Gomez, Hesham M. Abdullah, Yun Sun Lee, Danny J. Schnell, and Erich Grotewold. 2022. “Exploring Camelina Sativa Lipid Metabolism Regulation by Combining Gene Co-Expression and DNA Affinity Purification Analyses.” The Plant Journal n/a (n/a). https://doi.org/10.1111/tpj.15682.

Langfelder, Peter, and Steve Horvath. 2008. “WGCNA: An R Package for Weighted Correlation Network Analysis.” BMC Bioinformatics 9 (1): 559. https://doi.org/10.1186/1471-2105-9-559.

Langfelder, Peter, Bin Zhang, and Steve Horvath. 2008. “Defining Clusters from a Hierarchical Cluster Tree: The Dynamic Tree Cut Package for R.” Bioinformatics 24 (5): 719–20. https://doi.org/10.1093/bioinformatics/btm563.

Li, Chenxin, Joshua C. Wood, Anh Hai Vu, John P. Hamilton, Carlos Eduardo Rodriguez Lopez, Richard M. E. Payne, Delia Ayled Serna Guerrero, et al. 2022. “Single-Cell Multi-Omics Enabled Discovery of Alkaloid Biosynthetic Pathway Genes in the Medical Plant Catharanthus Roseus.” Preprint. Plant Biology. https://doi.org/10.1101/2022.07.04.498697.

Moghaddam, Samira Mafi, Atena Oladzad, Chushin Koh, Larissa Ramsay, John P. Hart, Sujan Mamidi, Genevieve Hoopes, et al. 2021. “The Tepary Bean Genome Provides Insight into Evolution and Domestication under Heat Stress.” Nature Communications 12 (1): 2638. https://doi.org/10.1038/s41467-021-22858-x.

Obayashi, T., and K. Kinoshita. 2009. “Rank of Correlation Coefficient as a Comparable Measure for Biological Significance of Gene Coexpression.” DNA Research 16 (5): 249–60. https://doi.org/10.1093/dnares/dsp016.

Shinozaki, Yoshihito, Philippe Nicolas, Noe Fernandez-Pozo, Qiyue Ma, Daniel J. Evanich, Yanna Shi, Yimin Xu, et al. 2018. “High-Resolution Spatiotemporal Transcriptome Mapping of Tomato Fruit Development and Ripening.” Nature Communications 9 (1): 364. https://doi.org/10.1038/s41467-017-02782-9.

Tippmann, S. 2015. “Programming Tools: Adventures with R.” Nature, no. 517: 109–10. https://doi.org/doi:10.1038/517109a.

Traag, V. A., L. Waltman, and N. J. van Eck. 2019. “From Louvain to Leiden: Guaranteeing Well-Connected Communities.” Scientific Reports 9 (1): 5233. https://doi.org/10.1038/s41598-019-41695-z.

Trapnell, Cole, Adam Roberts, Loyal Goff, Geo Pertea, Daehwan Kim, David R Kelley, Harold Pimentel, Steven L Salzberg, John L Rinn, and Lior Pachter. 2012. “Differential Gene and Transcript Expression Analysis of RNA-Seq Experiments with TopHat and Cufflinks.” Nature Protocols 7 (3): 562–78. https://doi.org/10.1038/nprot.2012.016.

Wickham, Hadley, Mara Averick, Jennifer Bryan, Winston Chang, Lucy McGowan, Romain François, Garrett Grolemund, et al. 2019. “Welcome to the Tidyverse.” Journal of Open Source Software 4 (43): 1686. https://doi.org/10.21105/joss.01686.

Wisecaver, Jennifer H., Alexander T. Borowsky, Vered Tzin, Georg Jander, Daniel J. Kliebenstein, and Antonis Rokas. 2017. “A Global Coexpression Network Approach for Connecting Genes to Specialized Metabolic Pathways in Plants.” The Plant Cell 29 (5): 944–59. https://doi.org/10.1105/tpc.17.00009.

